# Polyreactivity of antibodies from different B cell subpopulations is determined by distinct sequence patterns of variable region

**DOI:** 10.1101/2023.07.22.550160

**Authors:** Maxime Lecerf, Robin V. Lacombe, Jordan D. Dimitrov

## Abstract

An antibody molecule that is able to bind to multiple distinct antigens is defined as polyreactive. In the present study we performed statistical analyses to assess sequence correlates of polyreactivity of >600 antibodies cloned from different B cell types of healthy humans. The data reveled a number of sequence patterns of variable regions of heavy and light immunoglobulin chains that determine polyreactivity. The most prominent identified patterns were increased number of basic amino acid residues, reduced frequency of acidic residues, increased number of aromatic and hydrophobic residues, as well as longer length of CDR L1. Importantly, our study revealed that antibodies isolated from different B cell population used distinct sequence patterns (or combinations of them) for polyreactive antigen binding. Furthermore, we combined the data from sequence analyses with molecular modeling of selected polyreactive antibodies, and demonstrate that human antibodies can use multiple pathways for achieving antigen binding promiscuity. These data reconcile some contradictions in literature regarding the determinants of antibody polyreactivity. Moreover, our study demonstrates that mechanism of polyreactivity of antibodies evolves during immune response and might be tailored to specific functional properties of different B cell compartments. Finally, these data can be of use for efforts in development and engineering of therapeutic antibodies.

## Introduction

An antibody (Ab) molecule that has the capacity to interact with multiple unrelated antigens is referred to as polyreactive (1–3). Polyreactive Abs are normal constituent of immune repertoires and have important functions. For example, they participate in first line of defense against pathogens (4–6) and contribute for establishing mutualistic equilibrium between host and microbiome (7–9). Notably, many of broadly neutralizing Abs against HIV-1 and influenza virus display antigen-binding polyreactivity (10–16). Polyreactive antibodies also participate in clearance of apoptotic cells and damaged macromolecules (17–19). Nevertheless, polyreactivity is considered a negative trait for therapeutic Abs (20). It can cause deterioration of pharmacokinetics of therapeutic Abs and poses risk of undesirable effects (20). Moreover, polyreactivity correlates with other liabilities of Abs, such as tendency for self-association (21, 22).

Since discovery of polyreactive Abs, research efforts have been dedicated to unravel its molecular basis. These efforts were boosted following identification of polyreactivity as developability risk for therapeutic Abs. Many features of variable (V) domains have been associated with polyreactivity but controversial results are often reported in the literature. Thus, polyreactive Abs were shown to contain an elevated number of positively charged amino acid residues in their CDR H3 (15, 23–28). The presence of patches of positive charges on the molecular surface of antigen binding site or unbalanced distribution of charges have also been associated with polyreactivity of Abs (29–33). Analyses of synthetic Ab libraries revealed that some hydrophobic and aromatic residues in CDR H3 could also contribute to polyreactivity (34). Conversely, a recent study of large repertoires of mouse and human Abs, found a tendency for neutrality, or absence of a particular predominance of charges or hydrophobicity, in antigen binding sites of polyreactive Abs (35). Polyreactivity was also associated with longer CDR H3 loop (23, 27, 36, 37) and a higher tendency of this loop to form β-sheets (38). Other works, however, failed to detect difference in the size of CDR H3 between polyreactive and monoreactive Abs (8, 15, 24, 35). A global trait of V domains that has been related with promiscuity is conformational dynamics (3, 39). Many studies demonstrated that paratopes of polyreactive Abs have increased conformational flexibility (40–48). Nonetheless, there are reports demonstrating that some Abs can display polyreactivity without substantial conformational dynamics (13, 35, 49).

The conflicting results about the role of specific sequence or molecular attributes of Ab polyreactivity imply that there is still incomplete understanding of how Ab molecules attain promiscuous antigen binding. Several reasons can explain the conflicting results in the literature. They may be result of use of different experimental assays for assessment of Ab polyreactivity, use of different statistical methods for analyses of data, use of limited sets of Abs, evaluation of polyreactivity of Abs from different species (human vs mouse) and assessment of polyreactivity of Ab repertoires biased by selection for a particular antigen specificity. They may also reflect existence of multiple mechanisms of polyreactivity.

Recently we implemented a sequence analyses approach where the polyreactivity of clinical-stage therapeutic Abs was correlated with frequency of each amino acid residue, type of residues, and some global features (length of CDRs, number of somatic mutations) of V domains (32, 50, 51). These data revealed specific sequence traits associated with natural or induced by pro-oxidative substances polyreactivity in therapeutic Abs. We anticipated application of the same approach for analyses of large human Ab repertoires would allow deciphering determinants of polyreactivity.

We used data sets from two recent studies (27, 37) and analyzed the sequence patterns that determine the polyreactivity in human Ab repertoires. Importantly, we applied the analyses to subcategories of Abs identified on the basis on the B cell type from which they were cloned. We also applied molecular modelling to predict the structure of top polyreactive Abs from each category. Our data revealed several unique sequence patterns determining the polyreactivity of human Abs. Notably polyreactivity of Abs from different B cell subpopulations was determined by distinct sequence patterns of V domains. Our study also show that human Ab might use various molecular mechanisms of polyreactivity. These results might have important repercussions about understanding the mechanism and biological functions of polyreactive Abs.

## Results

### Human monoclonal Abs subjected to analyses

We used the data sets from two studies where monoclonal Abs were cloned from different subpopulations of healthy human B cells. Thus, we analyzed sequences of 398 Abs characterized in the study of Shehata et al (27). These Abs were cloned from naïve-, IgM memory-, and IgG memory B cells as well as from long-lived plasma cells (LLPC). In addition, we evaluated sequence correlates of polyreactivity of 240 monoclonal Abs cloned from IgA memory B cells, described in the study of Prigent et al (37). In both studies Abs were cloned without selection for a particular antigenic specificity and expressed exclusively as IgG1 for functional analyses. Of note, the two studies used different approaches for estimation of Ab polyreactivity. Thus, Shehata et al used technique referred to as polyspecificity reagent assay (PSR), where Ab reactivity to proteins available in human cell lysate is measured by flow cytometry and presented as gradually increasing numeric score (27). Prigent et al assessed Ab polyreactivity by ELISA (37). To use the semi-quantitative data presented in the work of Prigent et al. in our correlation analyses, we first assigned numeric scores of reactivity based on the number of recognized antigens by a given Ab (see Methods).

In previous works, V regions sequence patterns that determined polyreactivity were evaluated by using pools of Abs that originate from different B cell subpopulations. Moreover, some of these studies used Abs directed to pathogens (influenza virus or HIV-1) or Abs designed to be used as therapeutics. Here, we aimed to assess the sequence traits of V_H_ and V_L_ regions that correlate with Ab polyreactivity using Abs without pre-defined antigen specificity. We also considered important, in addition to bulk analyses, to segregate the Abs as based on their origin (B cell types from which they were isolated) and perform the statistical analyses individually.

### Correlation analyses of sequence features of VH and VL regions determining the polyreactivity of Abs from different B cell subpopulations

We applied Spearman non-parametric correlation analysis to assess the sequence patterns of V_H_ and V_L_ domains associated with polyreactivity of human Abs. The obtained data from correlation analysis for V_H_ domain are depicted on Figure 1. These data demonstrated that Ab polyreactivity significantly correlates with increased or reduced frequencies of certain amino acid residues in V domain. Strikingly, distinct patterns of significant correlations were observed between the groups of Abs originating from different B cells subpopulations (**Fig. 1**). Thus, polyreactivity negatively correlated with the number of V_H_ somatic mutations for Abs originating from IgM+ B cells, but no significant correlation was found in other groups. In addition, a positive correlation between the length of CDR H3 and polyreactivity was observed only for Abs cloned from IgA+ B cells. The polyreactivity of Abs from three B cells subtypes – naïve, IgG+ memory and plasma cells, correlated with the presence of a higher number of basic amino acid residues (Arg, His, Lys) in CDR H2 (naïve B cells and LLPC) and CDR H3 (IgG+ memory B cells) and within entire V_H_ region (LLPC). However, no increased number of basic amino acid residues in any part of V region was associated with polyreactivity of Abs isolated from IgM+ and IgA+ memory B cells. Reduced number of acidic amino acid residues (Asp and Glu) in entire V_H_ region or in CDR H1 and CDR H2 was associated with a higher polyreactivity in case of Abs cloned from all types of memory B cells but not from naïve B cells and LLPC (**Fig. 1**). Abs cloned from each B cell subpopulation demonstrated unique patterns of positive or negative correlates of polyreactivity with the number of polar amino acid residues in V_H_ domain. Presence of polar residues was especially important for determining polyreactivity of IgG+ memory B cells (**Fig. 1**). Thus, higher number of polar amino acids (as physiochemical group) in CDR H1 and H2 and in entire V_H_ was positively correlated with promiscuous antigen binding of Abs. Statistically significant correlations were also found for the number of Gln residues in entire V_H_ and Cys residues in CDR H3 (**Fig. 1**). Contrary, polyreactivity negatively correlated with the presence of Thr in CDR H1 in Abs cloned from naïve B cells. Our data demonstrated that polyreactivity of Abs isolated from IgA+ memory B cells significantly correlated with the presence of different hydrophobic residues (Val, Ile, Leu and Met) in entire V_H_ region or in CDR H1 and H3 loops (**Fig. 1**). This prominent pattern was not observed in Abs isolated from other types of B cells. On the contrary, the polyreactivity of Abs from IgG+ memory B cells was negatively correlated with the number of hydrophobic amino acids (as physiochemical group) in entire V_H_ domain or in CDR H2. The increased number of hydrophobic residues in V_H_ of Abs isolated from IgA+ B cells resulted in a significant correlation of polyreactivity with augmented hydrophobicity index (GRAVY) of all CDR loops and entire V_H_ domain (**Fig. 1**). Notable patterns of sequence correlations were also observed regarding the presence of aromatic amino acid residues in V_H_ domain. Thus, the number of aromatic amino acids (as a group) or of Tyr in V_H_ or in CDR H3 positively correlated with enhanced polyreactivity of Abs cloned from IgM+ B cells (**Fig. 1**). Positive correlation between polyreactivity and aromatic amino acids was also found for Abs from LLPC, but in this case the presence of any aromatic residues in CDR H1 or of Phe in CDR H3 reached significance. Notably, reverse tendencies were observed for Abs isolated from IgG+ and IgA+ B cells, where the presence of aromatic residues or Phe negatively correlated with the antigen binding polyreactivity (**Fig. 1**). The number of Gly residues significantly correlated with polyreactivity only for Abs that originated from naïve B cells. Thus, lower number of Gly in V_H_ region and CDR H2 was associated with higher polyreactivity. However, a reverse tendency was observed for Gly residues in case of CDR H1 (**Fig. 1**).

**Figure 1.**
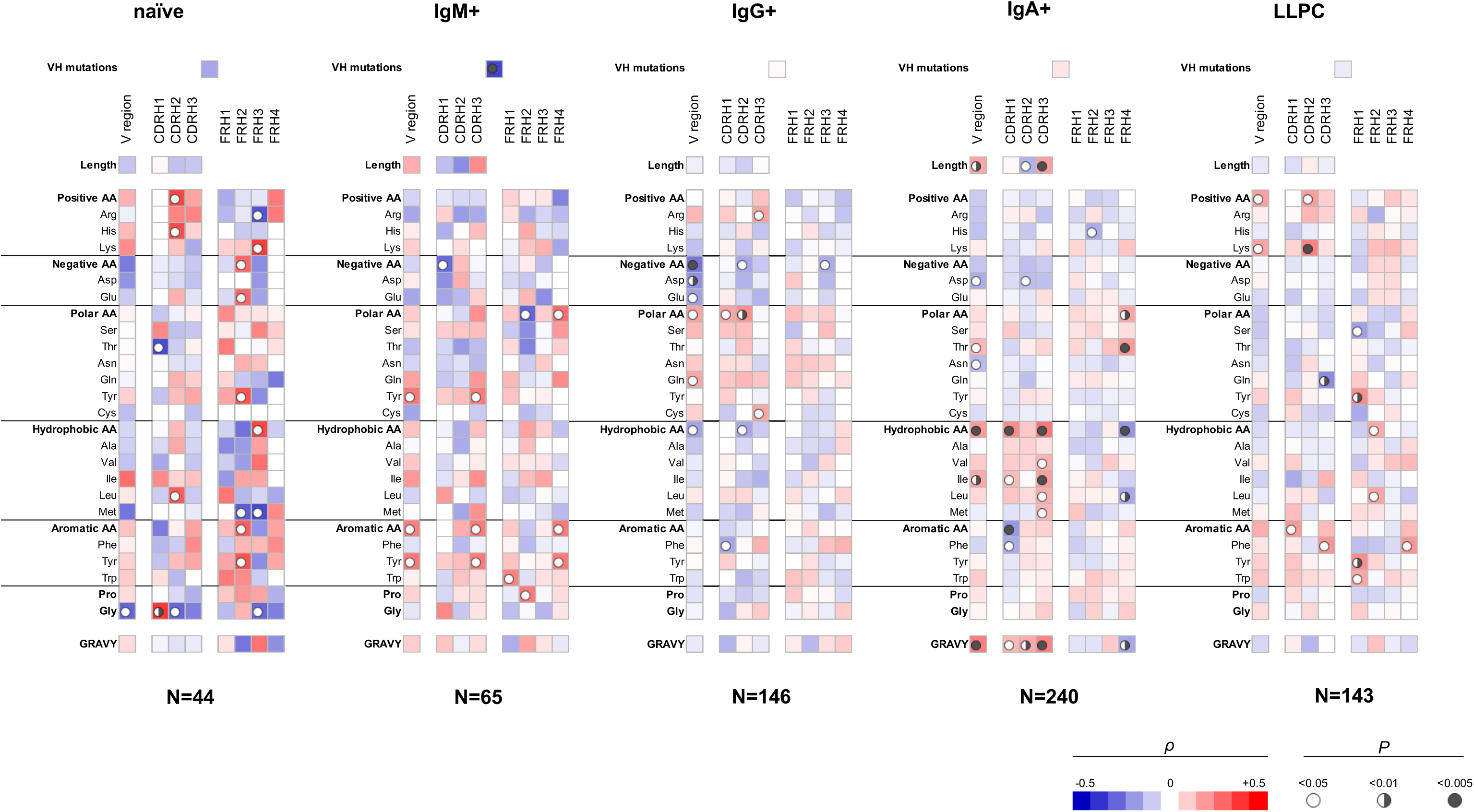
Correlation analyses of polyreactivity of human Abs cloned from different B cell subpopulation with sequence characteristics of V_H_ domain. The heat-maps present correlation coefficient (π) obtained by Spearman’s rank analyses of sequences characteristics of V_H_ domains (e.g., frequency of individual amino acid residues, type of residues, length of CDR H loops, mutation loads, hydrophobicity GRAVY index) with the extend of the antigen binding polyreactivity. Abs were grouped according to their B cell origin naïve; IgM+ memory; IgG+ memory, IgA+ memory, and long-lived plasma cells. The blue color signifies negative correlation, the red color signifies positive correlation. The statistical significance (p value) is indicated by circle. Open circle p < 0.05; semi-closed circle p < 0.01, and closed circle p < 0.005.

Our data also revealed that sequence features of framework regions (FW) of V_H_ can also contribute to polyreactivity of Abs. Importantly, we found that the patterns of significance in FW regions also depended of origin of Abs (**Fig. 1**). As FW regions are less frequently subjected to somatic mutations, the observed differences may reflect biased use of V_H_ genes in case of Abs with extended antigen binding polyreactivity.

Our analyses showed that a number of sequence features of V_L_ domain also correlate with polyreactivity of Abs (**Fig. 2**). Again, different sequence patterns of correlation were detected depending on the B cell origin of the studied Abs, albeit these differences were less prominent as compared to those observed in V_H_ (**Fig. 1** and **2**). An important observation from these data is the presence of a positive correlation between the size of entire V_L_ region or CDR L1 and polyreactive antigen binding in Abs cloned from three different B cell subpopulations, naïve, IgG+ memory and LLPC. Although, the data did not reach statistical significance same tendency was present in the case of IgM+ memory B cells. These data suggests that the length of CDR L1 is important determinant of polyreactivity of Abs. Interestingly, statistical significance was present between the lower numbers of Gln residues in CDR L1 and antigen binding polyreactivity in all B cell subpopulations except in naïve B cells (**Fig. 2**). Of note, this CDR L1 is characterized with highest variability in length among CDRs in V_L_ region. Another common sequence trait positively correlating with polyreactivity of different B cell subpopulations is the presence of a higher number of aromatic amino acids in all CDR loops. Such correlation reached significance for Abs cloned from naïve-, IgG+ and LLPC (**Fig. 2**). An important general observation of the correlation analyses of V_L_ domain is that the region that has the most significant capacity to determine the promiscuous antigen binding of Abs, from different B cell subpopulations is CDR L1 (**Fig. 2**). Similarly, as the observation for V_H_, the sequence features in FW regions correlating with polyreactivity of Abs showed particularities as dependent on the B cell type from which Abs were cloned.

**Figure 2.**
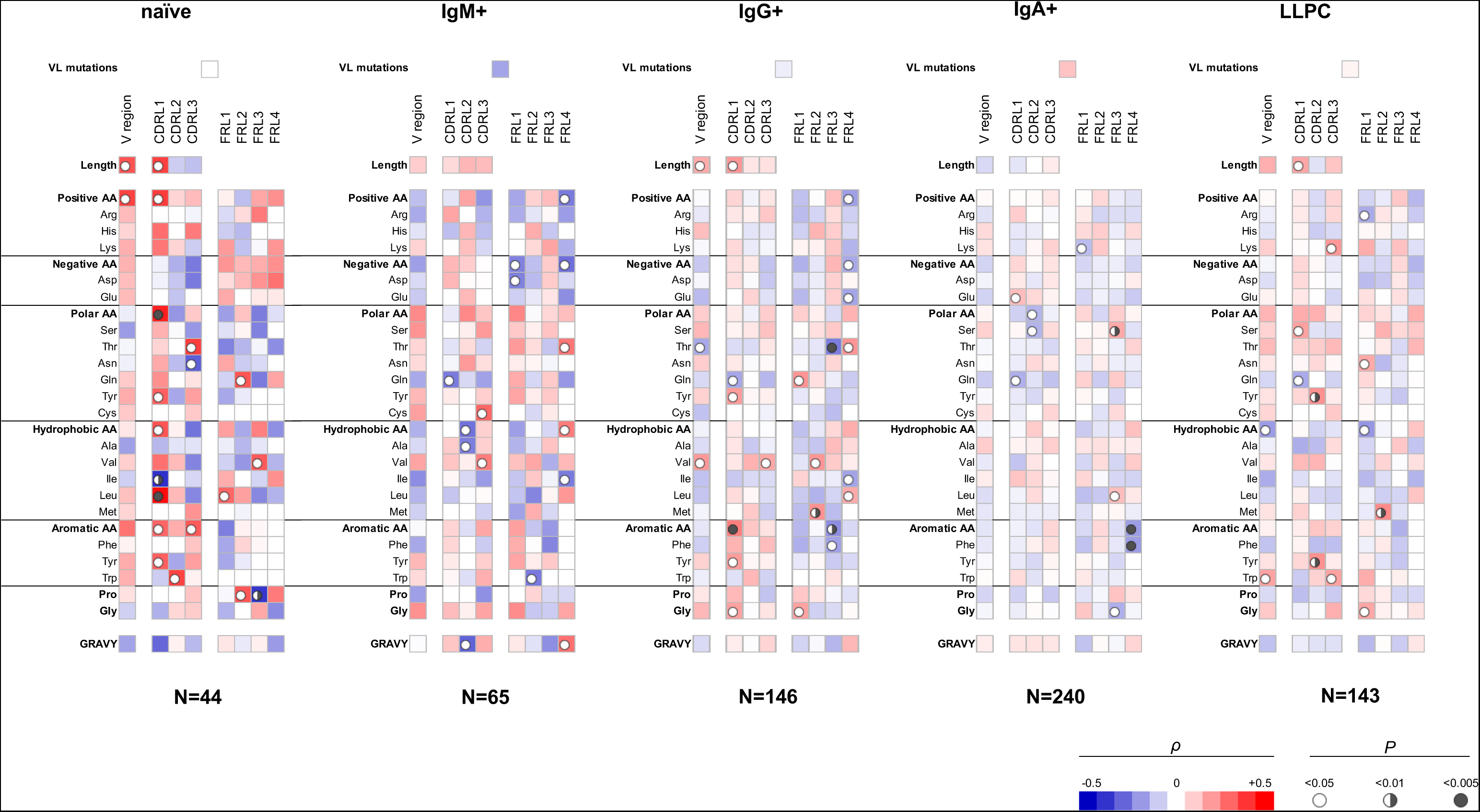
Correlation analyses of polyreactivity of human Abs cloned from different B cell subpopulation with sequence characteristics of V_L_ domain. The heat-maps present correlation coefficient (π) obtained by Spearman’s rank analyses of sequences characteristics of V_L_ domains (e.g., frequency of individual amino acid residues, type of residues, length of CDR L loops, mutation loads, hydrophobicity GRAVY index) with the extend of the antigen binding polyreactivity. Abs were grouped according to their B cell origin naïve; IgM+ memory; IgG+ memory, IgA+ memory, and long-lived plasma cells. The blue color signifies negative correlation, the red color signifies positive correlation. The statistical significance (p value) is indicated by circle. Open circle p < 0.05; semi-closed circle p < 0.01, and closed circle p < 0.005.

Taken together, the data from Spearman correlation analyses of V_H_ and V_L_ performed on Abs categorized as based on their B cell origin, unravel numerous sequence patterns significantly correlating with the antigen binding promiscuity. Notably, unique patterns were present in different groups of Abs.

### Compatibility between Ab repertoires

It is noteworthy that, the Abs cloned from IgA+ memory B cells (n=240) were analyzed for polyreactivity by ELISA (13), whereas the binding promiscuity of the rest of Abs (n=398) were assessed by PSR assay (27). We realized that the use of two distinct approaches can introduce bias in data and direct comparison of the results should be interpreted with a caution. However, previous analyses demonstrated that PSR and ELISA methods significantly correlate in their capacity to identify polyreactive Abs (21, 22). Validity of our results was also supported by the fact that there were considerable differences in the sequence patterns of V_H_ determining polyreactivity in groups of Abs that were assessed by an identical polyreactivity assay (PSR), as for example note the differences between Abs from IgG+ memory B cell group (n=146) and Abs from LLPC (n=143) (**Fig. 1**).

### Correlation analyses of sequence features of V_H_ and V_L_ regions determining the polyreactivity of Abs performed on integrated Ab repertoire

Next, we elucidated the sequence correlates of V regions that determine antigen binding polyreactivity in an integrated set of Abs originating from different B cell subpopulations. We excluded from this bulk repertoire only Abs originating from IgA+ B cells, as their polyreactivity was assessed by a different technique. By performing the statistical analyses of a panel of 398 Abs, a number of significant correlations were observed (**Fig. 3**). Thus, Ab polyreactivity in bulk repertoire correlated with the presence of higher number of Arg residues in CDR H3 and lower number acidic amino acids (especially Glu) in the entire V_H_ domain. Polyreactivity was also negatively correlated with Thr in CDR H1 and positively associated with the number of Gln residues in the whole V_H_. No significant correlation was observed between the number of hydrophobic amino acids in CDR loops of V_H_ domain and the polyreactivity of Abs. Notably, higher polyreactivity of Abs correlated with an increased number of aromatic residues (Phe and Trp) in CDR H3 (**Fig. 3**). The number of aromatic residues in FR1 was also significantly elevated in Abs with higher antigen binding polyreactivity.

**Figure 3.**
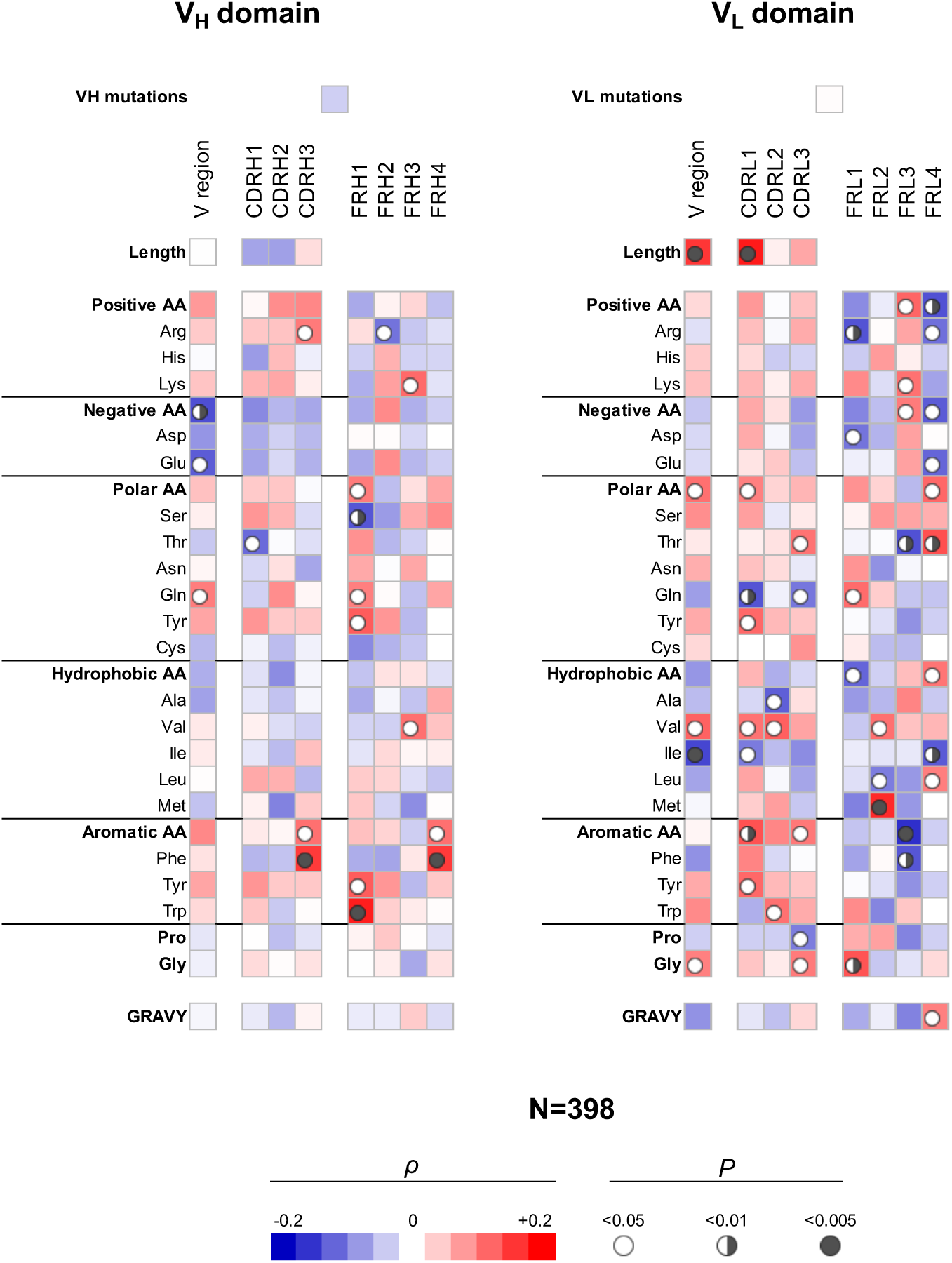
Correlation analyses of polyreactivity of human Abs cloned from different B cell subpopulation with sequence characteristics of V_H_ and V^L^ domains. The heat-maps present correlation coefficient (π) obtained by Spearman’s rank analyses of sequences characteristics of V_H_ (left panel) and V_L_ (right panel) domains with the extend of the antigen binding polyreactivity. Analyses were performed on integrated Ab repertoire consisting of Abs isolated from naïve; IgM+ memory; IgG+ memory, and long-lived plasma cells. The blue color signifies negative correlation, the red color signifies positive correlation. The statistical significance (p value) is indicated by circle. Open circle p < 0.05; semi-closed circle p < 0.01, and closed circle p < 0.005.

The correlation analyses revealed even a larger number of significant correlations of V_L_ sequence characteristics with the polyreactivity in bulk Ab repertoire (**Fig. 3**). Thus, a positive correlation was observed between the length of V_L_ region and particularly of CDR L1 and promiscuous antigen binding. Also, the number of polar amino acids in V_L_ or in CDR L1 and L3 correlated positively (Tyr and Thr) or negatively (Gln) with the Ab polyreactivity. The presence of the hydrophobic amino acid Val in entire V_L_ as well as in CDR L1 and L2 was similarly significantly associated with increased polyreactivity of Abs. However, the presence of Ile in whole V_L_ and Ala in CDR L2 had significantly negative impact on the Ab promiscuity (**Fig. 3**). Similarly, as in the case of V_H_, polyreactivity of Abs in integrated Ab repertoire positively correlated with the presence of aromatic amino acid residues in CDR L1 (Tyr), CDR L2 (Trp) and in CDR L3 (as type). Polyreactivity of Abs also significantly correlated with an increased number of Gly residues in V_L_ as well as in CDR L3 (**Fig. 3**).

Our analyses demonstrated that FW regions of light chain V regions have more decisive role for determining polyreactivity in integrated Ab repertoire as compared with the FW region of heavy V regions (**Fig. 3**). This observation suggests that larger set of V_L_ genes may be associated with the polyreactivity of Abs.

Collectively, these reseults revealed a number of significant correlates of Ab polyreactivity in bulk immune repertoire comprising Abs from distinct B cells subpopulations. These data showed certain prominent sequence patterns associated with polyreactivity – lack of acidic residues in V_H_, presence of aromatic amino acids in CDRs of both V_H_ and V_L_. The sequence pattern correlating with polyreactivity in bulk repertoire differed from the patterns in Abs from particular B cell subtype. Moreover, the strength of correlations was diminished, suggesting the presence of multiple pathways for achievement of polyreactive antigen binding.

### Analyses of Abs manifesting the highest level of antigen binding polyreactivity

To obtain further information about molecular correlates of polyreactivity, we next focused our analyses only on Abs that demonstrate the most prominent antigen binding promiscuity. To this end we selected top 5 polyreactive Abs from each B cell subpopulation. On Table 1 is presented summary of the characteristics of the variable regions of the Abs. These data revealed that the most polyreactive Abs did not have any specific bias in the lengths of CDR loops or their isoelectric points. Thus, CDR H3 and CDR L1 regions displayed the highest variability of lengths among different polyreactive Abs i.e., 8-26 and 6-12 residues for CDR H3 and CDR L1, respectively (Table 1). Theoretically calculated isoelectric points of CDRs also varied in very broad range between different Ab molecules (Table 1). For example, in the case of most diverse and important for antigen recognition region, CDR H3, the calculated pI values of the strongly polyreactive Abs were in the range between 3.67 and 9.14. It is noteworthy, that substantial variability in the length of CDR loops and the pI values were observed between Abs within all groups, except in case of Abs isolated from IgA+ B cells (Table 1). These data indicate that substantial level of antigen binding promiscuity can be achieved by alternative topological and physicochemical features of the antigen binding sites.

| Origin / B cell subset | Antibody | Polyreactivity score* | VH mutation number | CDRH1 |  | CDRH2 |  | CDRH3 |  | VL mutation number | CDRL1 |  | CDRL2 |  | CDRL3 |  |
| --- | --- | --- | --- | --- | --- | --- | --- | --- | --- | --- | --- | --- | --- | --- | --- | --- |
|  |  |  |  | Length (a.a.) | Isoelectric point | Length (a.a.) | Isoelectric point | Length (a.a.) | Isoelectric point |  | Length (a.a.) | Isoelectric point | Length (a.a.) | Isoelectric point | Length (a.a.) | Isoelectric point |
| Naive | ADI-45498 | 0,71069496 | 0 | 8 | 5.641 | 8 | 5.476 | 19 | 8.110 | 0 | 8 | 5.955 | 3 | 3.596 | 11 | 3.604 |
| Naive | ADI-45435 | 0,457177329 | 0 | 8 | 5.741 | 7 | 7.067 | 20 | 6.131 | 0 | 12 | 7.992 | 3 | 5.861 | 9 | 5.956 |
| Naive | ADI-45433 | 0,364564497 | 0 | 8 | 5.708 | 8 | 6.054 | 17 | 7.065 | 0 | 11 | 7.141 | 3 | 5.697 | 7 | 7.105 |
| Naive | ADI-45430 | 0,260866421 | 0 | 9 | 5.744 | 7 | 7.040 | 14 | 4.090 | 0 | 12 | 7.992 | 3 | 5.861 | 9 | 5.937 |
| Naive | ADI-46713 | 0,19010429 | 0 | 8 | 5.620 | 8 | 5.646 | 12 | 7.850 | 0 | 7 | 5.760 | 3 | 5.811 | 9 | 5.866 |
| IgM memory | ADI-45499 | 0,710238077 | 0 | 8 | 5.620 | 8 | 6.054 | 23 | 7.360 | 0 | 7 | 5.760 | 3 | 5.811 | 10 | 5.892 |
| IgM memory | ADI-47273 | 0,263842045 | 3 | 8 | 5.834 | 7 | 5.745 | 14 | 3.780 | 6 | 6 | 5.815 | 3 | 8.073 | 9 | 5.942 |
| IgM memory | ADI-45487 | 0,248689858 | 0 | 8 | 5.810 | 8 | 3.544 | 16 | 4.038 | 1 | 7 | 5.760 | 3 | 5.811 | 9 | 5.867 |
| IgM memory | ADI-45497 | 0,212869931 | 0 | 8 | 5.741 | 7 | 7.067 | 22 | 4.640 | 0 | 6 | 5.650 | 3 | 5.885 | 9 | 8.247 |
| IgM memory | ADI-45495 | 0,151349728 | 0 | 8 | 5.760 | 7 | 5.745 | 12 | 8.137 | 8 | 9 | 3.644 | 3 | 3.606 | 10 | 8.165 |
| IgG memory | ADI-47258 | 0,309315889 | 11 | 8 | 7.151 | 8 | 3.746 | 19 | 4.247 | 9 | 12 | 7.981 | 3 | 5.861 | 9 | 8.210 |
| IgG memory | ADI-47301 | 0,276256988 | 6 | 8 | 8.143 | 8 | 5.651 | 12 | 8.214 | 2 | 9 | 3.644 | 3 | 3.635 | 10 | 5.915 |
| IgG memory | ADI-47300 | 0,237031082 | 7 | 8 | 7.142 | 7 | 5.793 | 19 | 9.139 | 4 | 6 | 5.714 | 3 | 8.073 | 9 | 5.930 |
| IgG memory | ADI-47253 | 0,23696223 | 8 | 8 | 5.761 | 8 | 3.598 | 14 | 4.091 | 9 | 6 | 5.719 | 3 | 5.885 | 9 | 5.880 |
| IgG memory | ADI-45432 | 0,222241241 | 16 | 8 | 3.612 | 7 | 3.625 | 26 | 4.521 | 5 | 12 | 7.992 | 3 | 5.861 | 9 | 5.929 |
| IgA memory | 1-208 | 4 | 15 | 8 | 5.753 | 8 | 5.683 | 15 | 6.117 | 13 | 6 | 4.089 | 3 | 5.885 | 9 | 6.044 |
| IgA memory | 1-211 | 4 | 15 | 8 | 3.497 | 7 | 5.868 | 12 | 8.101 | 15 | 6 | 6.905 | 3 | 5.775 | 9 | 8.269 |
| IgA memory | 1-235 | 4 | 7 | 8 | 5.820 | 8 | 3.617 | 14 | 6.297 | 6 | 6 | 8.268 | 3 | 5.885 | 9 | 5.781 |
| IgA memory | 1-239 | 4 | 12 | 8 | 5.716 | 8 | 5.879 | 8 | 8.311 | 4 | 6 | 5.721 | 3 | 5.811 | 9 | 3.651 |
| IgA memory | 1-253 | 4 | 13 | 8 | 5.817 | 8 | 5.886 | 14 | 6.110 | 3 | 7 | 5.674 | 3 | 5.885 | 10 | 5.920 |
| LLPCs | ADI-47276 | 0,567042536 | 27 | 8 | 5.873 | 8 | 6.259 | 22 | 7.917 | 11 | 6 | 5.807 | 3 | 8.073 | 9 | 7.123 |
| LLPCs | ADI-47265 | 0,487816941 | 16 | 8 | 5.834 | 8 | 5.958 | 23 | 7.808 | 5 | 7 | 5.663 | 3 | 5.811 | 10 | 5.947 |
| LLPCs | ADI-47275 | 0,4180419 | 24 | 8 | 5.815 | 8 | 5.802 | 11 | 4.041 | 3 | 12 | 8.062 | 3 | 5.861 | 9 | 8.295 |
| LLPCs | ADI-47090 | 0,24495424 | 15 | 8 | 4.042 | 8 | 8.256 | 18 | 3.670 | 7 | 8 | 3.559 | 3 | 3.569 | 11 | 3.626 |
| LLPCs | ADI-47303 | 0,201395637 | 18 | 8 | 3.581 | 8 | 8.618 | 12 | 7.112 | 10 | 7 | 8.574 | 3 | 5.774 | 9 | 3.609 |

\*For antibodies cloned from naive-, IgM+ memory-, IgG+ memory B cells and LLPC PSR scores are displayed as published in XXX. For IgA+ memory B cells the polyreactivity was assessed by ELISA and Hep-assays. A numeric score 0-4 was assigned based on the number of recognized antigens and strength of the signal.

Further, to gain more detailed information about the topology of the antigen binding sites of polyreactive Abs, we applied structural modelling algorithm Rosie-2 (Rosetta Commons). The side views of the most probable structures of V regions of Abs are shown on Figure 4. These models demonstrated that the highly polyreactive human Abs have diverse topologies of their variable regions. One apparent feature observed in a part of the Abs is the presence of long and protruding CDR H3 (**Fig. 4**). This structural feature was especially evident in polyreactive Abs from naïve-B cells (ADI-45498), IgM+ memory B cells (ADI-45499, ADI-45497) and LLPC (ADI-47276 and ADI-47265) and well corresponded to the length of CDR H3 (Table 1). Interestingly, the structural models also indicated that some Abs have projecting only CDR L1 (ADI-45435, ADI-45430, ADI-47275) or in some cases both CDR H3 and CDR L1 are protruding (ADI-45435). Despite this obvious structural feature of variable regions of some of the polyreactive Abs, the structural models also showed that another fraction of highly polyreactive Abs did not display abnormally protruding CDR loops (**Fig. 4**). Indeed, the topology of their binding sites is similar as the one depicted for most of protein binding Abs available in Ab structural databases. This observation further substantiated our results from correlation analyses, suggesting that polyreactive Abs use diverse pathways for achievement of broad antigen binding promiscuity.

**Figure 4.**
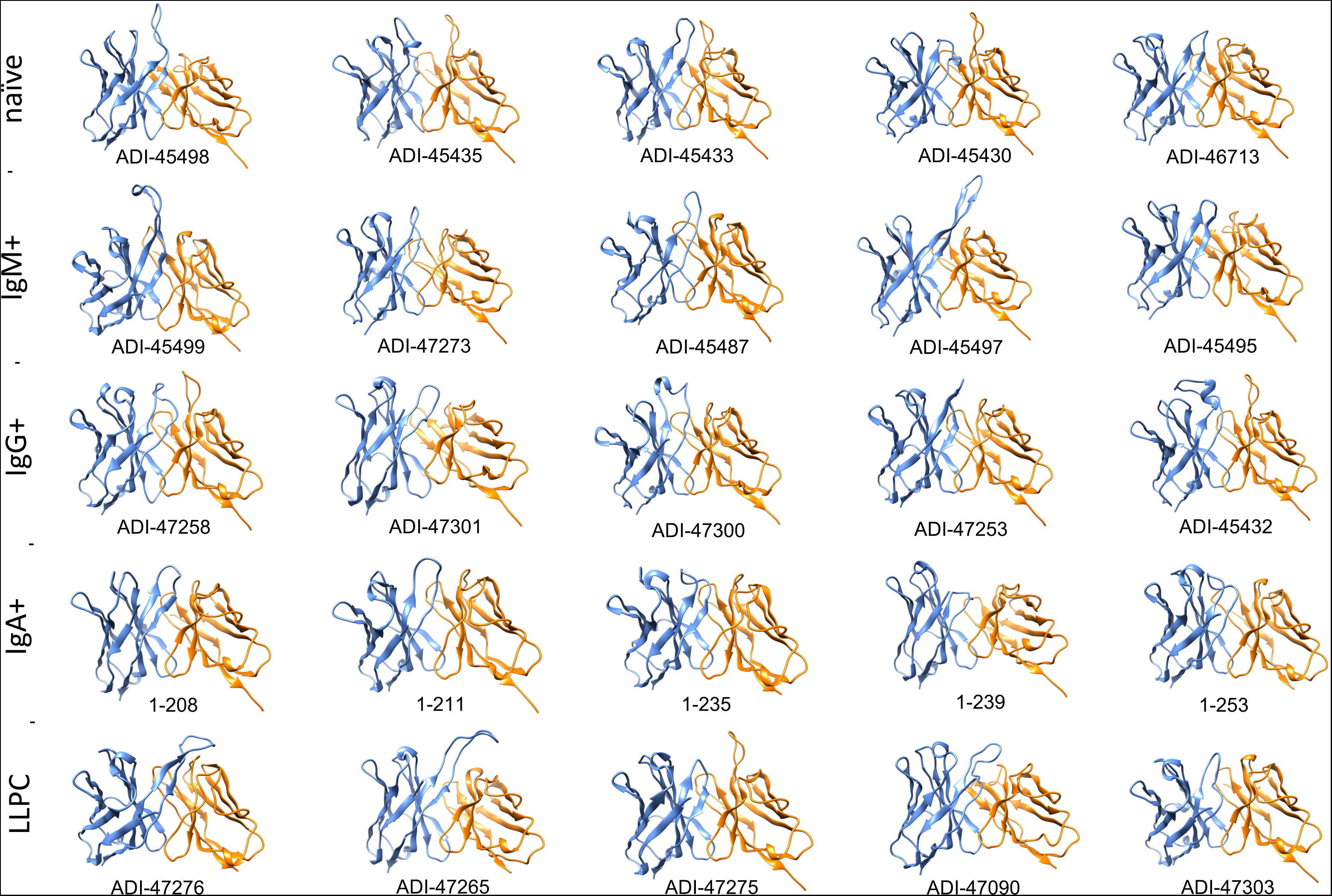
Molecular models of V regions of polyreactive Abs. Molecular models of the Abs manifesting the highest level of antigen binding polyreactivity from each B cell category. The Abs were ordered according the level of polyreactivity (PSR score, except for IgA+ memory B cells group) in a descending order from left to right. Side-view of the V region is presented. The blue ribbon corresponds to V_H_, the orange ribbon corresponds to V_L_. The models were generated with Rosie-2 Ab-module of Rosetta online server, and visualized by molecular viewer UCSF Chimera software v. 1.16.

Polyreactivity have been associated with specific physiochemical characteristics of antigen binding sites of Abs such as presence of patches of positive charges on the molecular surface. To elucidate whether this is the case in the human Abs isolated from different B cell compartments, we depicted the distribution of charges on the surface of antigen binding sites of selected set of Abs characterized with outstanding polyreactivity (**Fig. 5**). We also calculated three-dimensional Coulombic surface electrostatic potential of variable regions of the selected Abs (**Fig. 6**). These data demonstrated that different polyreactive Abs display distinct patterns of distribution of charged or non-polar residues in their binding sites (**Fig. 5**) and that the energy and space distribution of their electrostatic potentials differ substantially (**Fig. 6**). Some of the most polyreactive Abs, indeed, demonstrated extensive positively charged antigen binding sites (for example ADI-45498, ADI-46713, ADI-45499, ADI-47301, ADI-47300, ADI-47303), moreover the data revealed that a predominant fraction of Abs had antigen binding surfaces deprived of negative charges. Nonetheless, there were highly polyreactive Abs with balanced positive and negative charged patches or in some cases Abs had antigen binding site that carry little charged residues, or even predominantly negative charges (**Fig. 5 and 6**).

**Figure 5.**
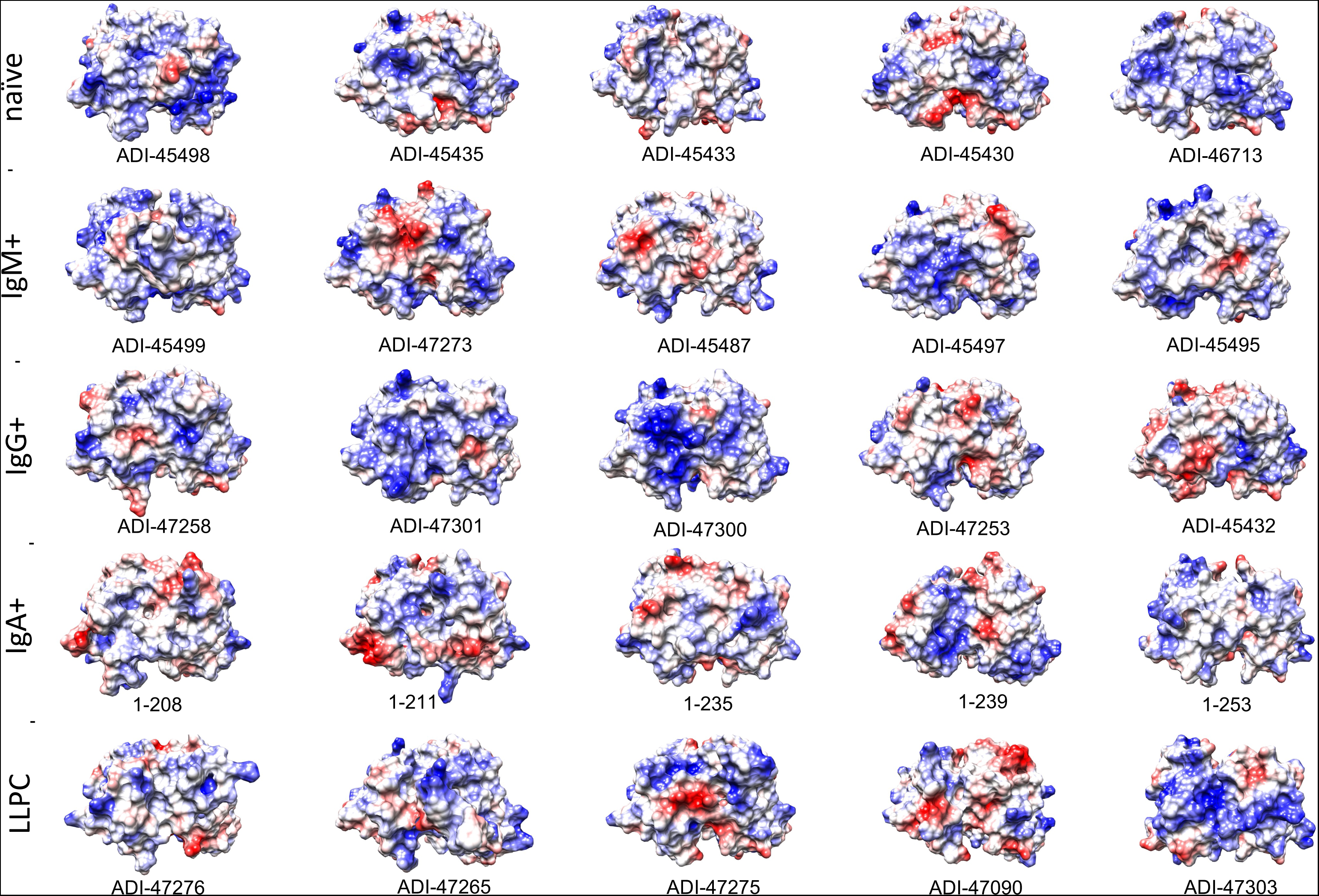
Surface electrostatic coloring of the antigen binding sites of polyreactive Abs. Top view of the structures of V regions of the most polyreactive Abs, isolated from different B cell subtypes. The V regions are ordered in descending order of their polyreactivity (from left to right). The structural models were generated by using Rosie-2 Ab-module of Rosetta online server. Electrostatic charges (blue: positive, red: negative) were depicted by using UCSF Chimera software v. 1.16.

**Figure 6.**
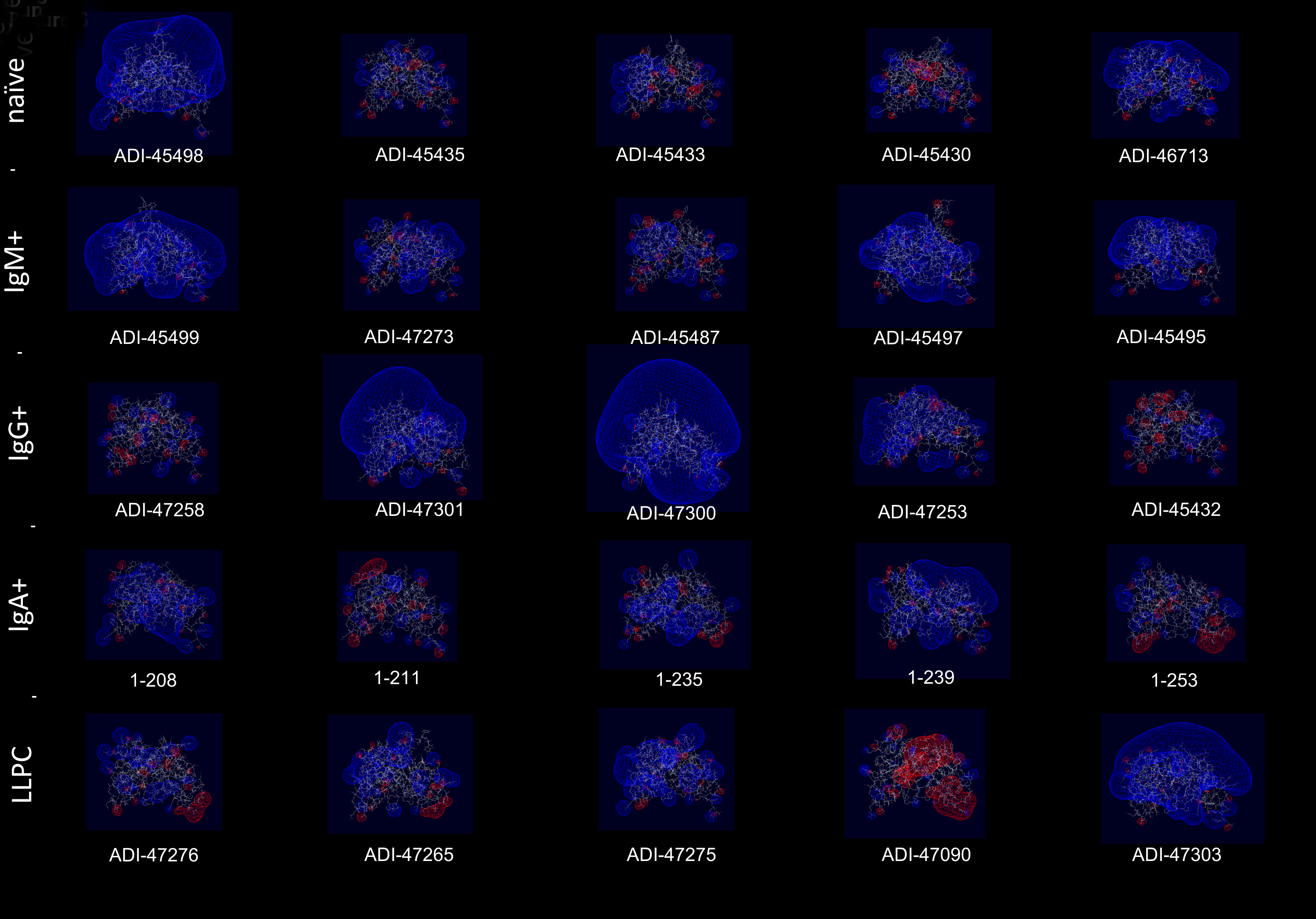
Three-dimensional electrostatic potential of V region of selected polyreactive Abs. Side view of the structural models of V regions of the most polyreactive Abs, isolated from different B cell subtypes. The models were obtained by using Rosie-2 Ab-module of Rosetta online server. The V regions are ordered in descending order of their polyreactivity (from left to right). The pictures show Coulombic electrostatic potentials (blue: positive, red: negative) that were calculated and visualized by using Swiss-PdbViewer v. 4.1.

Collectively, the data from this part of the study strongly suggest that the selected most prominently polyreactive Abs do not use universal topological or physiochemical qualities that determines the binding to multiple unrelated antigens. Rather variable regions with diverse properties can endow Abs with binding promiscuity.

## Discussion

In this study we deciphered the sequence correlates in V_H_ and V_L_ regions determining polyreactivity of human Abs. Our data show that Abs originating from different B cell subtypes relay on distinct sequence patterns for their promiscuous antigen binding. This finding suggests that mechanism of promiscuous antigen binding might evolve during the immune response, and it might be tailored to specific physiological roles of diverse B cell types. Our data confirmed some previously documented sequence determinants of polyreactivity, but they also highlighted several unrecognized ones. Furthermore, data from molecular modelling of V regions of selected Abs with the highest level of polyreactivity implied existence of alternative mechanisms that govern antigen binding promiscuity. Our data might explain some contradictions in the literature regarding the role of specific sequence or molecular features of V regions for polyreactivity.

To assess sequence traits of V domains associated with polyreactivity, we applied non-parametric correlation analyses (Spearman’s rank order correlation) using sequence and antigen binding data for >600 human Abs, unbiased by selection for a particular antigen specificity. The advantage of this approach is that it evaluates the individual contribution for Ab polyreactivity of every amino acid residue (or type of residues) in all parts of V domain (CDRs and FWs). As polyreactivity/monoreactivity of Abs is not binary measure but it rather spreads as a continuum of varying extents, the correlation analyses can reveal subtle details in the sequence determinants of antigen-binding promiscuity without the need for a priori grouping of Abs as polyreactive and monoreactive. A key aspect of this study is that we applied the correlation analyses not only to bulk Ab repertoires, but also to groups of Abs stratified based on B cell type from which they originate: naïve, IgM+ memory, IgG+ memory, IgA+ memory and long-lived plasma cells.

Our data showed that Abs originating from various human B cell subpopulations have different sequence determinants of their polyreactivity, especially in their V_H_ domain (Fig. 1 and 2). From analyzing the heatmaps depicted on Fig. 1, we can conclude that Abs relay on five major groups of sequence attributes for their polyreactivity – i) an increased number of positively charged residues in CDRs; ii) a reduced number of negatively charge residues in entire V_H_ and CDRs; iii) an increased number of aromatic residues in V_H_ and CDRs; iv) a reduced number of aromatic residues in CDR H1; v) an increased number of polar residues in V_H_ and CDRs, and vi) an increased number of hydrophobic residues in V_H_ and CDRs. Notably, our data indicated that different categories of Abs use unique combination of these sequence features for achieving polyreactivity and in some cases polyreactivity correlates with contrasting among the groups sequence characteristics. For example, polyreactive Abs cloned from naïve B cells relay on higher number of positively charged residues in their CDR H2. Similarly, higher polyreactivity of Abs cloned from LLPC correlated with the presence of an increased number of positively charged residues in CDRs. However, the polyreactivity in this group of Abs is also associated with a significantly elevated number of aromatic residues in CDR H3. On the other hand, polyreactivity of Abs from IgG+ memory B cells correlated strongly with reduced numbers of negatively charged residues, increased number of Arg in CDR H3 and an increased frequency of polar amino acid residues in V_H_ and CDRs. In contrast to other categories, Abs from IgM+ memory B cells relay exclusively on aromatic residues for promiscuous antigen binding. Interestingly, a unique pattern was detected also for Abs from IgA+ memory B cells (Fig. 1). Thus, polyreactivity of these Abs significantly correlated with the presence of a higher number of hydrophobic amino acid residues in V_H_ and CDRs (Leu, Ile, Met) and an increased length of CDR H3. The polyreactivity of Abs isolated from IgA+ B cells correlates with significantly reduced number of aromatic residues in CDR and tendency for lower prevalence of positively charged residues.

The V_L_ sequence characteristics that had strong correlation with polyreactivity was the length of CDR L1 and the presence of aromatic residues in CDRs. Of note, these patterns were constantly present in V_L_ domains of Abs from all groups with exception of Abs cloned from IgM+ and IgA+ memory B cells (Fig. 2). The elevated frequency of positively charged amino acid residues in V_L_ significantly correlated with polyreactivity only in case of Abs cloned from naïve B cells. The CDR L1 region displays the highest variability in size among the three CDR L loops. One can speculate that the longer CDR loops can contribute for higher conformational dynamics, thus promoting promiscuous antigen binding.

In addition to identified determinants in CDRs, the statistical analyses uncovered important correlations of different amino acids in FWs of V_H_ and V_L_ with polyreactivity. These correlations were especially pronounced in both V domains of Abs isolated from naïve- and IgM+ memory B cells. The association of sequence traits of FWs in unmutated or weakly mutated V regions with polyreactivity might indicate biased usage of specific genes encoding V domains. Indeed, contribution of some variable gene segments for polyreactivity has already been demonstrated in different Ab repertoires (15, 27, 35).

A sequence attribute that has been associated with polyreactivity of Abs is the mutation load in V_H_ and V_L_ domains. However, in the literature there is controversial evidence about the role of this attribute. Thus, some studies show that percentage of polyreactive Abs is the highest among germ-line Abs, whereas other works demonstrated higher prevalence of polyreactivity among Abs with mutated V regions (15, 23, 24, 27, 37, 41, 52, 53). Our analyses revealed that, that low number of mutations in V_H_ significantly correlates with polyreactivity of Abs cloned from IgM+ memory B cells (Fig. 1). In contrast, the mutations in V_H_ or V_L_ do not show significant correlation with binding promiscuity of human Abs.

Previous works demonstrated association of higher number of positively charged amino acid residues in CDRs or presence of positively charged patches with polyreactivity of Abs (15, 23, 26–33). Our statistical analyses confirmed these reports, but they also clearly demonstrated that this is not the determinant of polyreactivity for all groups of Abs and that this is not the strongest polyreactivity determinant. Molecular modelling of electrostatic potentials (Fig. 5 and 6) of top polyreactivity also demonstrated that positive charges are not exclusively present in all V regions. The statistical analyses demonstrated that there are two additional drivers of polyreactivity – increased prevalence of aromatic amino acids in CDRs of V_H_ and V_L_, as well as an increased hydrophobicity of CDRs. Our analyses, suggest that the former is the strongest correlate for polyreactivity present both in V_H_ and V_L_. In the literature, there are not systemic observation linking the higher number of aromatic amino acid or hydrophobic amino acids (as in the case of positively charged residues) as drivers of polyreactivity. There, are however, sporadic reports showing association of Trp, Val and Phe with polyreactivity in artificial Ab libraries (34) as well as in case of some engineered monoclonal broadly neutralizing HIV-1 Abs, where introduction of single or few aromatic residues resulted in a dramatic gain of antigen binding promiscuity (13, 54).

As consequence of their specific physicochemical and steric properties aromatic residues in CDR would provide capacity of Abs to establish numerous interactions with different macromolecules (55–57). High level of hydrophobic patches also gives a particular pattern of promiscuity through binding with similar hydrophobic regions on other macromolecules (57).

The finding that Abs originating from different B cell subpopulations have different sequence features that determines their polyreactivity might have important biological repercussions. These mechanisms may reflect different functional requirements of binding promiscuity for Abs from different B cell subpopulations. Thus, the fact that Abs cloned from IgA+ memory B cells use mostly hydrophobic residues as determinants of polyreactivity may reflects that these Abs operate mainly at mucosal surfaces. Indeed, the important role of Ab promiscuity of mucosal IgA has been emphasized by various recent works (7, 8, 36, 58, 59). We hypothesize that polyreactivity that relay on hydrophobicity is required for function of mucosal IgA, as intestinal and respiratory epithelial surfaces are covered with mucous layers that is strongly negatively charged (60). If polyreactivity of IgA was mediated by positively charged residues or patches in V regions as in other cases, these Abs would be restrained by binding to highly charged mucous and would not be able to interact with bacteria or other pathogens. Hydrophobicity of IgA may also be directing Abs for binding to hydrophobic motifs on bacterial cell wall, preventing bacterial adhesion (61). On the other hand, predominant role of aromatic residues for polyreactivity of IgM+ memory B cells may offer capacity of these Abs to bind to repetitive polysaccharide epitopes, that are typical for T cell-independent antigens. Indeed, the aromatic amino acid are the main type of amino acids implicated in recognition of glycans (62).

The length of CDR loops has been related to polyreactivity of antibodies (23, 27, 36, 37). Indeed, longer CDR would result in higher probability for conformational flexibility and as a consequence adaptability to various epitopes that may result in autoreactivity and polyreactivity (63, 64). The visualization of V regions of the most polyreactive Abs, indeed demonstrated that some molecules has outstandingly longer CDR loops, especially CDR H3 and CDR L1 (Table 1 and Fig. 4). However, many of the modelled Abs did not show longer than average CDRs. This result strongly suggests that albeit some of polyreactive antibodies may use long CDRs and eventual extended structural dynamics of binding site for achieving polyreactivity, this mechanism is not employed by all Abs able to bind multiple antigens.

In conclusion, this study provides evidence for implications of different molecular mechanisms in polyreactivity of human Abs. It also demonstrates that the sequence features that determine polyreactivity of Abs depend strongly on B cell origin of Abs. Thus, the study contributes valuable information about polyreactivity during evolution of immune response. Moreover, it reveals a number of important correlates of antigen binding promiscuity that would be of use for assessment of developability or engineering of therapeutic Abs.

## Methods

### Antibody repertoires and correlation analyses

We used data sets from two studies (27, 37) where polyreactivity of human monoclonal Abs was assessed. The study of Shehata et published the sequence information of all monoclonal Abs. The sequence information for Abs in the study of Prigent et al. was kindly provided by Dr Hugo Mouquet (Institute Pasteur, Paris, France). Since the polyreactivity in the study of Prigent et al was displayed in semi-quantitative value we assigned numerical values as based on the number of recognized antigens from ELISA panel Since the polyreactivity in the study of Prigent et al was displayed in semi-quotative quantitative value we assigned a polyreactivity score as based on the number of antigens for which each antibody was considered as reactive by authors (score ranking from 0 to 4 antigens)

The identification of CDR regions and assessment of the number of amino acid replacements in V domains was performed by IMGT/V-Quest alignment software. (https://www.imgt.org/). GRAVY index was determined with GRAVY Calculator (https://www.gravy-calculator.de) The polyreactivity of Abs was correlated with different features of variable regions – number of somatic mutations in V_H_ and V_L_ domains, lengths of CDR loops, number of charged, polar, aromatic, and hydrophobic residues in the CDR and FW regions, and frequency of individual amino acid residues in CDR and FW regions. The correlation analyses were performed by non-parametric Spearman’s rank-order analysis using Graph Pad Prism v.10 software (La Jolla, CA). A significant correlation was considered only this with *p* value ≤ 0.05.

### Modeling of V regions of top polyreactive Abs and calculation of Coulombic electrostatic potentials

For modeling structures of the V regions of selected Abs that manifested the highest level of antigen binding polyreactivity, we applied Ab structure modeling algorithm - RosettaAntibody3 (65–68). The sequences of V regions were submitted to Rossie-2 module of Rosetta online server (https://rosie.rosettacommons.org/). Relaxed three-dimensional models with lowest energy of coupled V_H_ and V_L_ domains were visualized by using Chimera UCSF Chimera package. The same software was used for coloring of the electrostatic charges of the molecular surfaces of the antibodies. Chimera is developed by the Resource for Biocomputing, Visualization, and Informatics at the University of California, San Francisco (supported by NIGMS P41-GM103311) (69). The three-dimensional Coulombic electrostatic potential was calculated and visualized by using Swiss-PdbViewer v. 4.1, http://www.expasy.org/spdbv/ (70).

## Acknowledgments

We are grateful to Dr Hugo Mouquet (Pasteur Institute, Paris, France) for providing us complete sequence information of antibodies cloned from IgA+ B cells and for fruitful discussions. This work was supported by Institut National de la Santé et de la Recherche Médicale (INSERM, France) and by grants from Agence Nationale de la Recherche (ANR-13-JCV1-006-01) and from the European Research Council (Project CoBABATI ERC-StG-678905), both to J.D.D.

## References

1. A. L. Notkins, Polyreactivity of antibody molecules. Trends Immunol 25, 174–179 (2004).

2. J. D. Dimitrov et al., Antibody polyreactivity in health and disease: statu variabilis. J Immunol 191, 993–999 (2013).

3. D. Jain, D. M. Salunke, Antibody specificity and promiscuity. Biochem J 476, 433–447 (2019).

4. M. Boes, A. P. Prodeus, T. Schmidt, M. C. Carroll, J. Chen, A critical role of natural immunoglobulin M in immediate defense against systemic bacterial infection. J Exp Med 188, 2381–2386 (1998).

5. A. F. Ochsenbein et al., Control of early viral and bacterial distribution and disease by natural antibodies. Science 286, 2156–2159 (1999).

6. Z. H. Zhou et al., The broad antibacterial activity of the natural antibody repertoire is due to polyreactive antibodies. Cell Host Microbe 1, 51–61 (2007).

7. F. Fransen et al., BALB/c and C57BL/6 Mice Differ in Polyreactive IgA Abundance, which Impacts the Generation of Antigen-Specific IgA and Microbiota Diversity. Immunity 43, 527–540 (2015).

8. J. J. Bunker et al., Natural polyreactive IgA antibodies coat the intestinal microbiota. Science 358 (2017).

9. J. W. Chen et al., Autoreactivity in naive human fetal B cells is associated with commensal bacteria recognition. Science 369, 320–325 (2020).

10. B. F. Haynes et al., Cardiolipin polyspecific autoreactivity in two broadly neutralizing HIV-1 antibodies. Science 308, 1906–1908 (2005).

11. H. Mouquet et al., Polyreactivity increases the apparent affinity of anti-HIV antibodies by heteroligation. Nature 467, 591–595 (2010).

12. M. Liu et al., Polyreactivity and autoreactivity among HIV-1 antibodies. J Virol 89, 784–798 (2015).

13. J. Prigent et al., Conformational Plasticity in Broadly Neutralizing HIV-1 Antibodies Triggers Polyreactivity. Cell Rep 23, 2568–2581 (2018).

14. G. Bajic et al., Autoreactivity profiles of influenza hemagglutinin broadly neutralizing antibodies. Sci Rep 9, 3492 (2019).

15. J. J. Guthmiller et al., Polyreactive Broadly Neutralizing B cells Are Selected to Provide Defense against Pandemic Threat Influenza Viruses. Immunity 53, 1230–1244 e1235 (2020).

16. A. Reyes-Ruiz, J. D. Dimitrov, How can polyreactive antibodies conquer rapidly evolving viruses? Trends Immunol 42, 654–657 (2021).

17. M. Y. Chou et al., Oxidation-specific epitopes are dominant targets of innate natural antibodies in mice and humans. J Clin Invest 119, 1335–1349 (2009).

18. M. R. Ehrenstein, C. A. Notley, The importance of natural IgM: scavenger, protector and regulator. Nat Rev Immunol 10, 778–786 (2010).

19. Z. H. Zhou et al., Polyreactive antibodies plus complement enhance the phagocytosis of cells made apoptotic by UV-light or HIV. Sci Rep 3, 2271 (2013).

20. O. Cunningham, M. Scott, Z. S. Zhou, W. J. J. Finlay, Polyreactivity and polyspecificity in therapeutic antibody development: risk factors for failure in preclinical and clinical development campaigns. MAbs 13, 1999195 (2021).

21. T. Jain et al., Biophysical properties of the clinical-stage antibody landscape. Proc Natl Acad Sci U S A 114, 944–949 (2017).

22. T. Jain, T. Boland, M. Vasquez, Identifying developability risks for clinical progression of antibodies using high-throughput in vitro and in silico approaches. MAbs 15, 2200540 (2023).

23. H. Wardemann et al., Predominant autoantibody production by early human B cell precursors. Science 301, 1374–1377 (2003).

24. T. Tiller et al., Autoreactivity in human IgG+ memory B cells. Immunity 26, 205–213 (2007).

25. S. Birtalan et al., The intrinsic contributions of tyrosine, serine, glycine and arginine to the affinity and specificity of antibodies. J Mol Biol 377, 1518–1528 (2008).

26. K. E. Tiller et al., Arginine mutations in antibody complementarity-determining regions display context-dependent affinity/specificity trade-offs. J Biol Chem 292, 16638–16652 (2017).

27. L. Shehata et al., Affinity Maturation Enhances Antibody Specificity but Compromises Conformational Stability. Cell Rep 28, 3300–3308 e3304 (2019).

28. Y. D. Kwon et al., Improved pharmacokinetics of HIV-neutralizing VRC01-class antibodies achieved by reduction of net positive charge on variable domain. MAbs 15, 2223350 (2023).

29. A. Datta-Mannan et al., Balancing charge in the complementarity-determining regions of humanized mAbs without affecting pI reduces non-specific binding and improves the pharmacokinetics. MAbs 7, 483–493 (2015).

30. A. Schoch et al., Charge-mediated influence of the antibody variable domain on FcRn-dependent pharmacokinetics. Proc Natl Acad Sci U S A 112, 5997–6002 (2015).

31. L. A. Rabia, Y. Zhang, S. D. Ludwig, M. C. Julian, P. M. Tessier, Net charge of antibody complementarity-determining regions is a key predictor of specificity. Protein Eng Des Sel 31, 409–418 (2018).

32. M. Lecerf, A. Kanyavuz, S. Lacroix-Desmazes, J. D. Dimitrov, Sequence features of variable region determining physicochemical properties and polyreactivity of therapeutic antibodies. Mol Immunol 112, 338–346 (2019).

33. M. I. J. Raybould et al., Five computational developability guidelines for therapeutic antibody profiling. Proc Natl Acad Sci U S A 116, 4025–4030 (2019).

34. R. L. Kelly, D. Le, J. Zhao, K. D. Wittrup, Reduction of Nonspecificity Motifs in Synthetic Antibody Libraries. J Mol Biol 430, 119–130 (2018).

35. C. T. Boughter et al., Biochemical patterns of antibody polyreactivity revealed through a bioinformatics-based analysis of CDR loops. Elife 9 (2020).

36. M. A. Berkowska et al., Circulating Human CD27-IgA+ Memory B Cells Recognize Bacteria with Polyreactive Igs. J Immunol 195, 1417–1426 (2015).

37. J. Prigent et al., Scarcity of autoreactive human blood IgA(+) memory B cells. Eur J Immunol 46, 2340–2351 (2016).

38. J. M. J. Laffy et al., Promiscuous antibodies characterised by their physico-chemical properties: From sequence to structure and back. Prog Biophys Mol Biol 128, 47–56 (2017).

39. D. Jaiswal, S. Verma, D. T. Nair, D. M. Salunke, Antibody multispecificity: A necessary evil? Mol Immunol 152, 153–161 (2022).

40. V. Manivel, N. C. Sahoo, D. M. Salunke, K. V. Rao, Maturation of an antibody response is governed by modulations in flexibility of the antigen-combining site. Immunity 13, 611–620 (2000).

41. V. Manivel, F. Bayiroglu, Z. Siddiqui, D. M. Salunke, K. V. Rao, The primary antibody repertoire represents a linked network of degenerate antigen specificities. J Immunol 169, 888–897 (2002).

42. J. Yin, A. E. t. Beuscher, S. E. Andryski, R. C. Stevens, P. G. Schultz, Structural plasticity and the evolution of antibody affinity and specificity. J Mol Biol 330, 651–656 (2003).

43. L. C. James, P. Roversi, D. S. Tawfik, Antibody multispecificity mediated by conformational diversity. Science 299, 1362–1367 (2003).

44. R. Jimenez, G. Salazar, K. K. Baldridge, F. E. Romesberg, Flexibility and molecular recognition in the immune system. Proc Natl Acad Sci U S A 100, 92–97 (2003).

45. H. P. Nguyen et al., Germline antibody recognition of distinct carbohydrate epitopes. Nat Struct Biol 10, 1019–1025 (2003).

46. T. Khan, D. M. Salunke, Adjustable locks and flexible keys: plasticity of epitope-paratope interactions in germline antibodies. J Immunol 192, 5398–5405 (2014).

47. M. L. Fernandez-Quintero et al., Characterizing the Diversity of the CDR-H3 Loop Conformational Ensembles in Relationship to Antibody Binding Properties. Front Immunol 9, 3065 (2018).

48. R. J. Blackler et al., Antigen binding by conformational selection in near-germline antibodies. J Biol Chem 298, 101901 (2022).

49. D. K. Sethi, A. Agarwal, V. Manivel, K. V. Rao, D. M. Salunke, Differential epitope positioning within the germline antibody paratope enhances promiscuity in the primary immune response. Immunity 24, 429–438 (2006).

50. M. Lecerf, A. Kanyavuz, S. Rossini, J. D. Dimitrov, Interaction of clinical-stage antibodies with heme predicts their physiochemical and binding qualities. Commun Biol 4, 391 (2021).

51. M. Lecerf, R. Lacombe, A. Kanyavuz, J. D. Dimitrov, Functional Changes of Therapeutic Antibodies upon Exposure to Pro-Oxidative Agents. Antibodies (Basel) 11 (2022).

52. Z. H. Zhou, A. G. Tzioufas, A. L. Notkins, Properties and function of polyreactive antibodies and polyreactive antigen-binding B cells. J Autoimmun 29, 219–228 (2007).

53. H. X. Liao et al., Initial antibodies binding to HIV-1 gp41 in acutely infected subjects are polyreactive and highly mutated. J Exp Med 208, 2237–2249 (2011).

54. R. Diskin et al., Restricting HIV-1 pathways for escape using rationally designed anti-HIV-1 antibodies. J Exp Med 210, 1235–1249 (2013).

55. F. L. Gervasio, R. Chelli, P. Procacci, V. Schettino, The nature of intermolecular interactions between aromatic amino acid residues. Proteins 48, 117–125 (2002).

56. E. J. Sundberg, R. A. Mariuzza, Molecular recognition in antibody-antigen complexes. Adv Protein Chem 61, 119–160 (2002).

57. H. Ausserwoger et al., Non-specificity as the sticky problem in therapeutic antibody development. Nat Rev Chem 6, 844–861 (2022).

58. J. J. Bunker, A. Bendelac, IgA Responses to Microbiota. Immunity 49, 211–224 (2018).

59. K. E. Huus, C. Petersen, B. B. Finlay, Diversity and dynamism of IgA-microbiota interactions. Nat Rev Immunol 21, 514–525 (2021).

60. J. Leal, H. D. C. Smyth, D. Ghosh, Physicochemical properties of mucus and their impact on transmucosal drug delivery. Int J Pharm 532, 555–572 (2017).

61. M. C. van Loosdrecht, J. Lyklema, W. Norde, G. Schraa, A. J. Zehnder, The role of bacterial cell wall hydrophobicity in adhesion. Appl Environ Microbiol 53, 1893–1897 (1987).

62. K. L. Hudson et al., Carbohydrate-Aromatic Interactions in Proteins. J Am Chem Soc 137, 15152–15160 (2015).

63. L. Yu, Y. Guan, Immunologic Basis for Long HCDR3s in Broadly Neutralizing Antibodies Against HIV-1. Front Immunol 5, 250 (2014).

64. W. K. Wong, J. Leem, C. M. Deane, Comparative Analysis of the CDR Loops of Antigen Receptors. Front Immunol 10, 2454 (2019).

65. A. Sivasubramanian, A. Sircar, S. Chaudhury, J. J. Gray, Toward high-resolution homology modeling of antibody Fv regions and application to antibody-antigen docking. Proteins 74, 497–514 (2009).

66. N. A. Marze, S. Lyskov, J. J. Gray, Improved prediction of antibody VL-VH orientation. Protein Eng Des Sel 29, 409–418 (2016).

67. B. D. Weitzner et al., Modeling and docking of antibody structures with Rosetta. Nature protocols 12, 401–416 (2017).

68. B. D. Weitzner, J. J. Gray, Accurate Structure Prediction of CDR H3 Loops Enabled by a Novel Structure-Based C-Terminal Constraint. J Immunol 198, 505–515 (2017).

69. E. F. Pettersen et al., UCSF Chimera--a visualization system for exploratory research and analysis. Journal of computational chemistry 25, 1605–1612 (2004).

70. N. Guex, M. C. Peitsch, SWISS-MODEL and the Swiss-PdbViewer: an environment for comparative protein modeling. Electrophoresis 18, 2714–2723 (1997).

